# Leg immobilization and subsequent recovery resistance training affect skeletal muscle angiogenesis related markers in young healthy adults regardless of prior resistance training experience

**DOI:** 10.1101/2024.11.24.625075

**Authors:** Mason C. McIntosh, J. Max Michel, Joshua S. Godwin, Daniel L. Plotkin, Derick A. Anglin, Madison L. Mattingly, Anthony Agyin-Birikorang, Nicholas J. Kontos, Harsimran S. Baweja, Matt S. Stock, C. Brooks Mobley, Michael D. Roberts

## Abstract

We recently reported that resistance trained (T, n=10) and untrained (UT, n=11) young adults experience vastus lateralis (VL) muscle atrophy following two weeks of disuse, and 8 weeks of recovery resistance training (RT) promotes VL hypertrophy in both participant cohorts. However, angiogenesis targets and muscle capillary number were not examined and currently no human studies that have sought to determine if disuse followed by recovery RT affects these outcomes. Thus, we examined whether disuse and/or recovery RT affected these outcomes. All participants underwent two weeks of left leg immobilization using locking leg braces and crutches followed by eight weeks (3d/week) of knee extensor focused progressive RT. VL biopsies were obtained at baseline (PRE), immediately after disuse (MID), and after RT (POST). Western blotting was used to assay angiogenesis markers and immunohistochemistry was performed in 16/21 participants to determine type I and II muscle fiber capillary number. Significant main effects of time (p<0.05) were observed for protein levels of VEGF (MID<POST), VEGFR2 (PRE&MID<POST), TSP-1 (PRE<POST), TIMP1 (MID<POST), phosphorylated/pan eNOS (Ser1177) (POST<PRE), and pan eNOS (PRE<POST). VEGFR2 exhibited a training status*time (p=0.018), but no differences existed between T and UT at any time point. A significant main effect of time was observed for type II fiber capillary number (PRE<POST), and type II fiber cross-sectional area (fCSA) increased from MID to POST (+25%, p<0.001) and PRE to POST (+20%, p=0.019). No significant correlations exist for percentage changes in type II fiber capillary number and type II fCSA from PRE-to-MID (r= 0.020), MID-to-POST (r= 0.392), or PRE-to-POST (r= −0.120) across all participants (p>0.100). Although disuse and recovery RT affect skeletal muscle angiogenesis-related protein targets, prior training history does not differentially affect these outcomes.

**NEW AND NOTEWORTHY:** This is the first study to examine how limb immobilization and recovery resistance training affect molecular outcomes related to angiogenesis in younger adults with or without a prior training history. Regardless of resistance training history, the molecular responses are largely similar between participant cohorts and is suggestive of a reduced (pre-mid) and increased (mid-post) angiogenic response, with disuse and subsequent recovery resistance training.

## INTRODUCTION

Skeletal muscle is a tissue that robustly adapts to various loading stimuli (1), and mechanical overload through resistance training (RT) promotes remodeling to support myofiber and whole tissue hypertrophy (2). Although the mobilization of satellite cells and anabolic signaling through the mammalian target of rapamycin complex 1 (mTORC1) signaling have been generally appreciated for promoting skeletal muscle hypertrophy (3), recent evidence indicates angiogenesis (or the process of capillary expansion or growth) may also play a significant role (4). To this end, longer-term RT promotes increases in skeletal muscle capillarization (5–7), lower pre-training muscle capillary density is associated with limited hypertrophic outcomes in adults (8–10), and genetic mouse models indicate that mechanical overload-induced skeletal muscle hypertrophy is blunted when angiogenesis is impaired (11, 12). From a mechanistic standpoint, RT in humans upregulates the mRNA and protein expression of vascular endothelial growth factor (VEGF) (13–15), a potent pro-angiogenic factor (4, 16). There is also evidence that RT can dynamically alter matrix metalloproteinases (MMPs) as well as their inhibitors (TIMPs) (17, 18), and these proteins are also involved in the extracellular matrix (ECM) remodeling needed for angiogenesis (19). Finally, limited data indicates that the angiogenesis response may be more robust in untrained (UT) individuals when compared to trained (T) individuals suggesting that diminished angiogenesis may be an adaptive response to RT (15).

Contrary to RT-induced skeletal muscle hypertrophy, disuse promotes remodeling processes that result in skeletal muscle tissue and myofiber atrophy (20). Disuse related skeletal muscle loss is a common outcome among medically injured patients, those with common illnesses, and aging individuals (20). Appreciable skeletal muscle atrophy can manifest within 10 days of disuse, equating in the approximate loss of ∼0.5-0.6% of muscle mass per day (21) and results in up to 30% loss of total quadriceps size after 90-120 days (22, 23). These maladaptive responses are, due to altered net protein balance where muscle protein synthesis does not outweigh breakdown, largely due to decreases in muscle protein synthesis (24). Additional mechanisms associated with disuse-induced skeletal muscle atrophy include a dysregulation in muscle-to-nerve communication, an increase in endoplasmic reticulum stress, and a loss in ribosome content (23–26). Disuse via unilateral leg immobilization has been reported to reduce skeletal muscle capillary content (27), and this may coincide with the atrophic process.

While disuse and mechanical overload potentially operate through different signaling pathways, the molecular response to skeletal muscle disuse and subsequent recovery RT is well-elucidated. Moreover, although RT can increase skeletal muscle capillarization, limited data indicates that the angiogenesis response to RT may be more robust in untrained (UT) individuals when compared to trained (T) individuals (15), which suggests that diminished angiogenesis may be an adaptive response to RT. Thus, examining the angiogenesis response to disuse and recovery RT in individuals with or without a prior history of resistance training would provide valuable insight.

We previously reported that T (n=10) and UT (n=11) young adults experienced similar magnitudes of muscle atrophy according to ultrasound measures following two weeks of disuse, and 8 weeks of recovery RT promoted skeletal muscle hypertrophy in both participant cohorts (28). However, given that angiogenesis markers were not originally assayed, we sought to determine if the disuse and recovery RT protocol affected several angiogenesis and vascular remodeling markers (VEGF, VEGFR2, TSP-1, MMP2, MMP9, TIMP1, TIMP2, total eNOS, and phosphorylated eNOS) or type I and II muscle fiber capillarization, and whether responses differed between UT and T participants. We developed multiple *a priori* hypotheses. First, disuse and recovery RT would downregulate and subsequently upregulate skeletal muscle protein levels of VEGF, VEGR2, MMP2, MMP9, and phosphorylated as well as total eNOS (angiogenesis markers) in both UT and T participants, albeit this response would be more robust in UT participants. Second, disuse and recovery RT would upregulate and subsequently downregulate skeletal muscle protein levels of angiogenesis inhibitors (i.e., TSP-1, TIMP1, TIMP2) in both UT and T participants. We additionally hypothesized that changes observed in the protein expression of these angiogenic regulators would be more robust in UT versus T. Finally, we hypothesized muscle capillary number would be greater in T versus UT participants throughout the intervention and that changes in muscle capillary number throughout the intervention would be significantly associated with changes in muscle fiber size.

## METHODS

### Ethical approval and participant screening

This study is a follow-up investigation of a study approved by the Auburn University Institutional Review Board and in accordance with the most recent revisions of *Declaration of Helsinki* (IRB protocol #23-220 MR2305, clinical trial registration # NCT05760066). The participants were males and females from the local area who met the following criteria: (i) 18-35 years old; (ii) no known cardiometabolic disease (e.g. diabetes, hypertension, heart disease) or any musculoskeletal condition contraindicating participation in exercise training or donating skeletal muscle biopsies; (iii) free from metal implants that would interfere with x-ray based data collection; (iv) had not consumed known anabolic agents that affect hormone status within the past two months (e.g. exogenous testosterone, growth hormone, etc.); (v) free from blood clotting disorders that would contraindicate donating a muscle biopsy; (vi) females were not pregnant or attempting to become pregnant. Additional participant information and protocol for participant recruitment and screening can be found in Michel et al. (28).

### Study design

A more detailed description of the study design and methodology can be found in Michel et al. (28). Briefly, UT (n=11, 26±3 years old, 78.4±24.4 kg, 8 males, 3 females) and T (n=10, 27±3 years old, 81.4 ± 8.1 kg, 8 males, 2 females) participants underwent 2 weeks of left leg immobilization via locking brace and crutches followed by 8 weeks of RT. Participants visited the laboratory for baseline data collection and to be fitted for their locking leg brace and crutches (PRE), after two weeks of disuse for subsequent data collection and removal of their locking brace and crutches (MID), and after 8 weeks of RT for final data collection (POST). All data collection sessions occurred a minimum of 72 hours after any RT session and the order of data collection events were held constant for each session.

### Leg disuse protocol

The leg disuse protocol was designed based on previous work by MacLennan et al. (29). Upon completion of all testing procedures during PRE, participants were fitted with a knee joint immobilizer brace (T Scope® Premier Post-Op Knee Brace; Breg Inc., Carlsbad, CA, USA). The brace was locked in place at ∼90° of knee flexion. In this position the participants’ knee extensors remained relaxed and unloaded to ensure the leg was fully non-weight bearing. Participants were additionally fitted with axillary crutches and were provided gait-training to effectively ambulate with a single leg while using crutches. After two weeks of bracing and ambulation on crutches, each participant returned to the laboratory for Post-brace testing. At this time the brace was removed, and each participant returned to their bilateral full weight bearing gait.

### Post-brace resistance training protocol

Following brace removal, participants underwent 8 weeks of supervised RT at a frequency of three (non-consecutive) days per week. Participants performed barbell back squats, leg extensors, barbell bench press, lat pulldown, and lying hamstring curls on day 1 and 3 of the week. On day 2 of training, the individuals performed leg press, hex-bar deadlift, barbell row, overhead press, and dumbbell biceps curl. Progressive overload was implemented throughout the training period, and additional training details on training can be found in Michel et al. (28).

### PRE, MID, and POST testing sessions

#### Urine specific gravity, height, and body mass

During testing session visits, participants reported to the laboratory following an overnight fast. Hydration status was assessed via a urine specific gravity test measured with a handheld refractometer (ATAGO; Bellevue, WA, USA) with the goal of ensuring adequate hydration (value ≤1.020) (30). Body mass and height were assessed with a digital scale (Seca 769, Hanover, MD, USA).

#### Vastus lateralis muscle biopsies

After ultrasound scans, skeletal muscle biopsy samples were collected from the left VL a 5 mm Bergstrom needle. Biopsies at MID and POST were taken ∼2 cm proximal from the preceding biopsy scar. Briefly, participants laid supine on an athletic training table and the upper thigh was shaved and cleaned with 70% isopropanol prior to receiving an injection of 1% lidocaine (0.6-0.9 mL). After waiting ∼5 minutes to allow lidocaine to take full effect, the area was cleaned with chlorhexidine and a pilot incision made with a sterile, single-use No. 11 surgical blade (AD Surgical, Sunnyvale, CA, USA). The biopsy was then collected using a 5 mm Bergstrom needle under suction. Approximately 50-100 mg of skeletal muscle tissue was collected, immediately teased of blood and connective tissue, and separated for histological and biochemical analysis. Tissue (∼30 mg) was mounted for histology at a 90° angle on a piece of cork in a 1:1 w/w mixture of optimal cutting temperature (OCT) solution and tragacanth powder (Alfa Aesar, Ward Hill, MA, USA). The mount was then covered in OCT, frozen in liquid nitrogen cooled 2-methylbutane for ∼30 seconds, then contained in a box top floating atop liquid nitrogen before long-term storage at −80°C. A separate ∼20-60 mg tissue sample was placed in foil and flash frozen in liquid nitrogen for protein isolation and western blotting. All tissue triage procedures occurred within a 2-minute window.

### Wet laboratory analyses

#### Western blotting

Muscle tissue (∼20 mg) was lysed using a general cell lysis buffer (Cell Signaling Technology, Danvers, MA, USA; Cat. No. 9803) and tight-fitting pestles. Lysates were then centrifuged at 500 g for 5 minutes and supernatants were placed into new 1.7 mL tubes. A commercially available BCA protein assay kit (Thermo Fisher, Waltham, MA, USA; Cat. No. A55864) and spectrophotometer (Agilent Biotek Synergy H1 hybrid reader; Agilent, Santa Clara, CA, USA) were used to determine supernatant protein concentrations. Thereafter, supernatants were prepared for western blotting at equal protein concentrations (1 µg/µL) using 4x Laemmli buffer and deionized water. Western blot preps (15 µL) were pipetted onto SDS gels (4-15% Criterion TGX Stain-free gels, Bio-Rad Laboratories; Hercules, CA, USA), and proteins were separated by electrophoresis at 180 V for 50 minutes. Proteins were then transferred to methanol-preactivated PVDF membranes (Bio-Rad Laboratories) for 2 hours at 200 mA. Following transfers, membranes were Ponceau stained for 10 minutes, washed with diH_2_O for ∼30 seconds, dried, and digitally imaged (ChemiDoc Touch, Bio-Rad). Following Ponceau imaging, membranes were reactivated in methanol, blocked with 5% non-fat bovine milk in tris-buffered saline with Tween-20 (TBST) for 1 hour, and washed 3x5 minutes in TBST. Membranes were then incubated with primary antibodies (1:1000 dilution in TBST containing 5% bovine serum albumin (BSA)) on a rocker overnight at 4°C (listed in Table 1).

**Table 1.** Antibodies used for western blotting and immunohistochemistry.

| Antibody | Host species (isotype) | Company (Cat. No.) |
| --- | --- | --- |
| *Western blotting* |  |  |
| VEGF | Rb | Cell Signaling (65373) |
| VEGFR2 | Rb | Cell Signaling (9698) |
| TSP-1 | Rb | Cell Signaling (37879) |
| MMP2 | Rb | Cell Signaling (40994) |
| MMP9 | Rb | Cell Signaling (13667) |
| TIMP1 | Rb | Cell Signaling (8946) |
| TIMP2 | Rb | Cell Signaling (5738) |
| eNOS | Rb | Cell Signaling (32027) |
| Phospho-eNOS (Ser1177) | Rb | Cell Signaling (9507) |
| Anti-Rb IgG, HRP-conjugated | G | Cell Signaling (7074) |
| *Immunohistochemistry* |  |  |
| CD31/PECAM-1 | M (IgG1) | DSHB (P2B1) |
| Type I myosin heavy chain | M (IgG2b) | DSHB (BA-D5) |
| Dystrophin | Rb (IgG) | Abcam (ab218198) |
| Anti-M IgG1, AF555-conjugated | G | Thermo Fisher (A-21127) |
| Anti-M IgG2b, AF488-conjugated | M | Thermo Fisher (A-21141) |
| Anti-Rb IgG, AF647-conjugated | Rb | Thermo Fisher (A-21245) |

Abbreviations: DSHB, Developmental Studies Hybridoma Bank (Iowa City, IA, USA); G, goat; M, mouse; Rb, rabbit. Other note: protein acronyms are defined in text.

After overnight primary antibody incubations, antibody solutions were decanted, and membranes were washed for 3x5 minutes in TBST. The membranes were then incubated at room temperature for 60 minutes in in TBST containing 5% BSA and a 1:2000 v/v dilution of HRP-conjugated antibody against the host species of the primary antibody (presented in Table 1). The secondary antibody solution was decanted, and membranes were washed for 3x5 minutes in TBST. The membranes were then developed in a gel documentation system (ChemiDoc Touch, Bio-Rad) with enhanced chemiluminescent reagent (Luminata Forte HRP substrate; Millipore Sigma, Burlington, MA, USA), and band densitometry was performed using associated software. For non-phosphorylated targets, target band densities were obtained and divided by Ponceau densitometry values and fold-change values were derived by dividing Ponceau-normalized band density values by the aggregate PRE mean value of the UT group for each target. Band density values for phosphorylated eNOS was divided by corresponding pan band density values (after being normalized to ponceau stains), and again fold-change values were derived by dividing these values by the aggregate PRE mean value of the of T participants.

#### Immunohistochemistry for capillarization quantification

Biopsy samples preserved with optimal cutting temperature media were sectioned at a thickness of 12 μm using a cryotome (Leica Biosystems; Buffalo Grove, IL, USA), adhered to positively charged glass slides (VWR; Radnor, PA, USA), and stored at −80℃ until being batch processed for immunohistochemical analyses to detect capillaries, fiber type, dystrophin, and nuclei. During batch-processing, sections were removed from −80°C storage, air-dried at room temperature for ≥2 hours, and fixed with acetone at −20°C for 5 minutes. Slides were then incubated with 3% hydrogen peroxide for 10 minutes at room temperature, followed by a 1-minute incubation with autofluorescence quenching reagent (TrueBlack, Cat. No. 23007; Biotium, Fremont, CA, USA) and blocked for 1 hour in 2.5% horse serum in phosphate-buffered saline (PBS) at room temperature. After blocking, slides were incubated overnight at 4°C with a primary antibody cocktail in PBS containing 2.5% horse serum and 1:100 v/v dilutions of anti-PECAM-1, anti-type I myosin heavy chain, and anti-dystrophin antibodies listed in Table 1. The following day, the sections were washed with PBS 3x5 minutes and incubated for 60 minutes in a secondary antibody cocktail containing 1:250 v/v dilutions of the fluorophore-conjugated secondary antibodies listed in Table 1. Slides were then stained with DAPI (1:10,000; 4’, 6-diamidino-2-phenylindole; Cat. No. D3571; Thermo Fisher Scientific) for 10 minutes at room temperature and mounted with glass coverslips using 1:1 PBS and glycerol as mounting medium. Sections were stored in the dark at 4°C until imaging was conducted. Multiple digital 20x images per participants’ time points were captured with a fluorescence microscope at (Zeiss Axio Imager.M2) and motorized stage. All areas selected for analyses were free of freeze-fracture artifacts. Fiber cross-sectional area was conducted in Michel et al. (28). Capillary contacts to type I or type II fibers were manually quantified using a tally counter.

#### Statistical analyses

Data were plotted and analyzed in GraphPad Prism (v10.2.2). Data were first checked for normality using Shapiro-Wilk tests. Only VEGF protein in T participants, MMP9 in UT, and TIMP1 in UT were non-normally distributed (p<0.05). Thus, we opted to proceed with parametric analyses given that most dependent variables were normally distributed. All outcome variables were examined using two-way (training status*time) repeated measures ANOVAs with, and Tukey’s post hoc tests used to decompose significant main effects and/or interactions. Select Pearson’s correlations were conducted on certain change scores from immunohistochemistry analysis as explained in the results section. Statistical significance was established throughout as *p*<0.05.

## RESULTS

### Angiogenesis markers

Figure 1 contains skeletal muscle protein levels of positive (VEGF, VEGFR2) and negative (TSP-1) regulators of angiogenesis. VEGF exhibited a main effect of time (p=0.006; Fig. 1a) where PRE and POST values were greater than MID (p≤0.011; Fig. 1a). VEGFR2 exhibited a main effect of time (p<0.001) and a training status*time interaction (p=0.018; Fig. 1b). Although PRE values of the trained participants trended higher than the untrained participants (p=0.083), values between training status groups were not significantly different at MID (p=0.694) or POST (p=0.127). Values at PRE and MID, however, were less than POST in both training status groups (p≤0.047). TSP-1 exhibited a main effect of time (p=0.040) and a training status*time interaction (p=0.038; Fig. 1c). Despite a significant interaction, values were not different between T and UT at any timepoint (p≥.0.079), while values at PRE were less than MID in both training status groups (p≤0.026).

**Figure 1.**
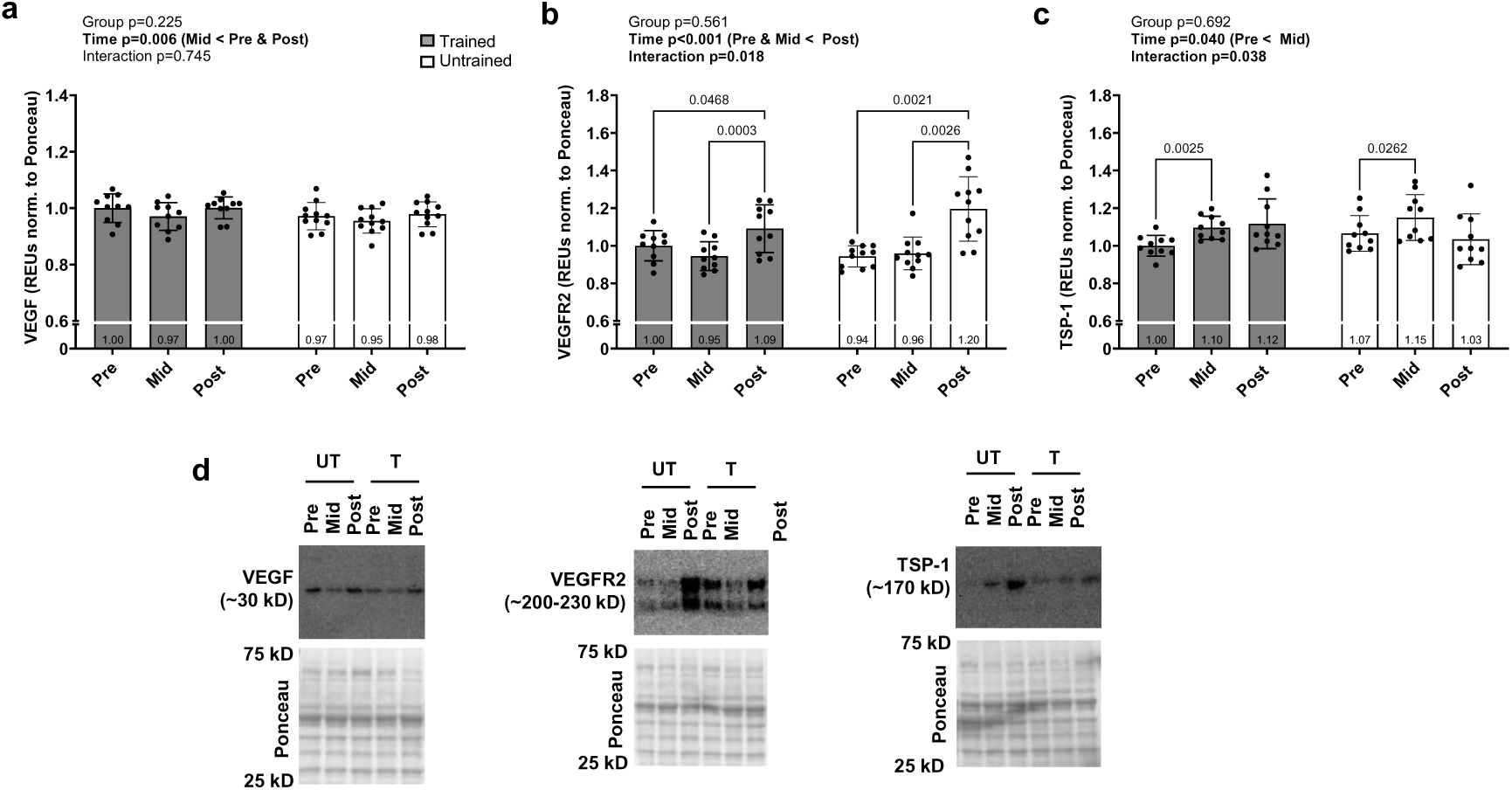
Levels of proteins associated with angiogenesis. Data presented as mean ± standard deviation for VEGF (a), VEGFR2 (b), and TSP-1 (c) prior to the intervention (Pre), following two weeks of leg disuse (Mid), and following 8 weeks of recovery resistance training (Post). Panel d shows representative western blots. Abbreviations: T, trained participant; UT, untrained participant. Other note: each participant’s data point is overlain on bar graphs and mean values are depicted at bottom of bar.

### Extracellular matrix remodeling markers

Figure 2 contains skeletal muscle protein expression data from select matrix metalloproteinases (MMPs) and MMP inhibitors (TIMP1/2) implicated in angiogenesis. Neither MMP2 (Fig. 2a), MMP9 (Fig. 2b), TIMP1 (Fig. 2c), nor TIMP2 (Fig. 2d) exhibited significant main or interaction effects.

**Figure 2.**
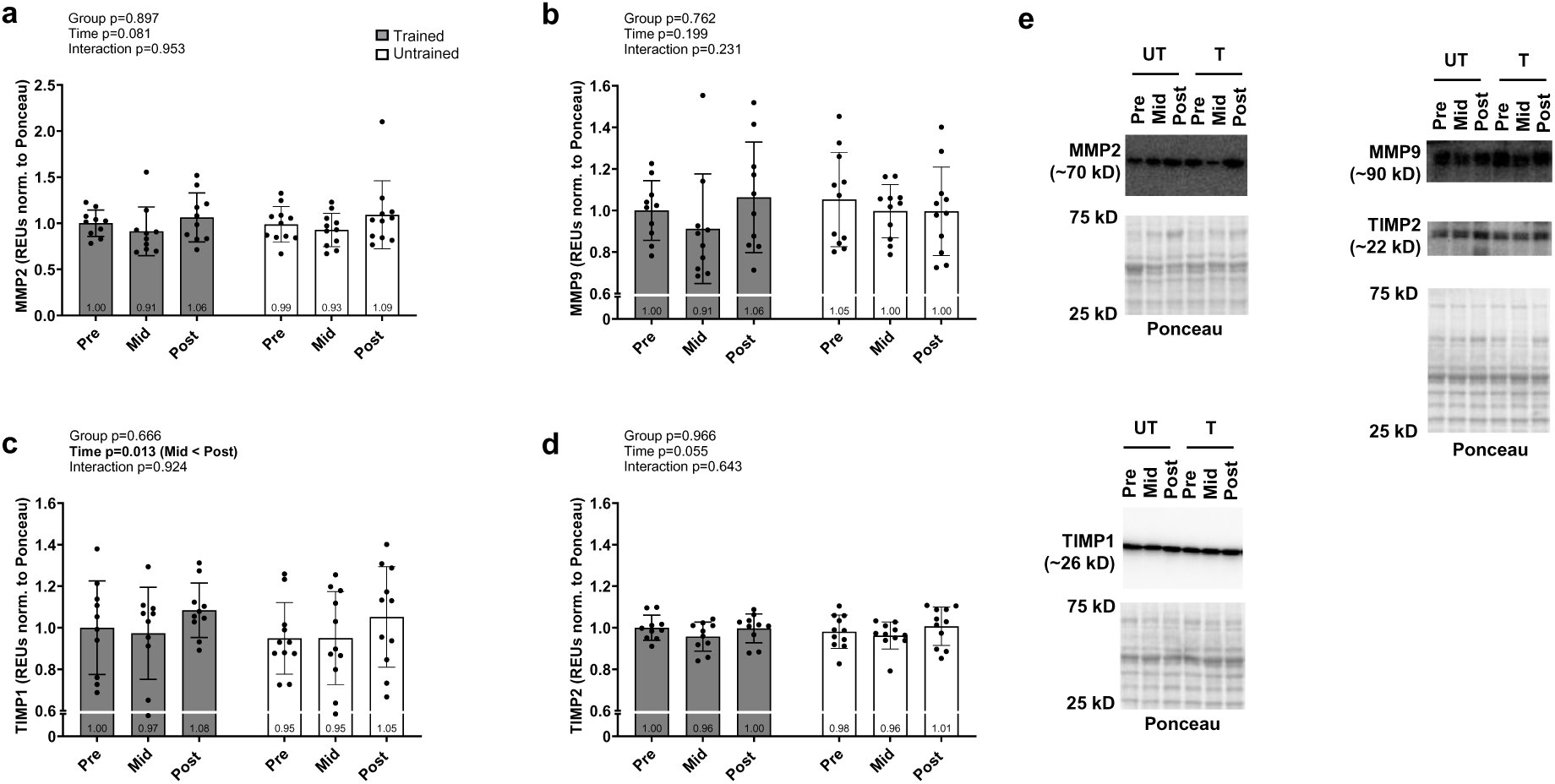
Levels of proteins associated with extracellular matrix remodeling during angiogenesis. Data presented as mean ± standard deviation for MMP2 (a), MMP9 (b), TIMP1 (c) and TIMP2 prior to the intervention (Pre), following two weeks of leg disuse (Mid), and following 8 weeks of recovery resistance training (Post). Panel e shows representative western blots. Abbreviations: T, trained participant; UT, untrained participant. Other note: each participant’s data point is overlain on bar graphs and mean values are depicted at bottom of bar.

### eNOS markers

Figure 3 contains skeletal muscle pan and phosphorylated eNOS levels. While significant interaction effects were not evident for these markers, significant main effects of time were evident for both. Specifically, phosphorylated eNOS levels were lower in all participants at POST versus PRE (p=0.017; Fig. 3a) whereas pan eNOS levels were greater at POST versus PRE (p=0.044) and MID (p=0.001; Fig. 3b).

**Figure 3.**
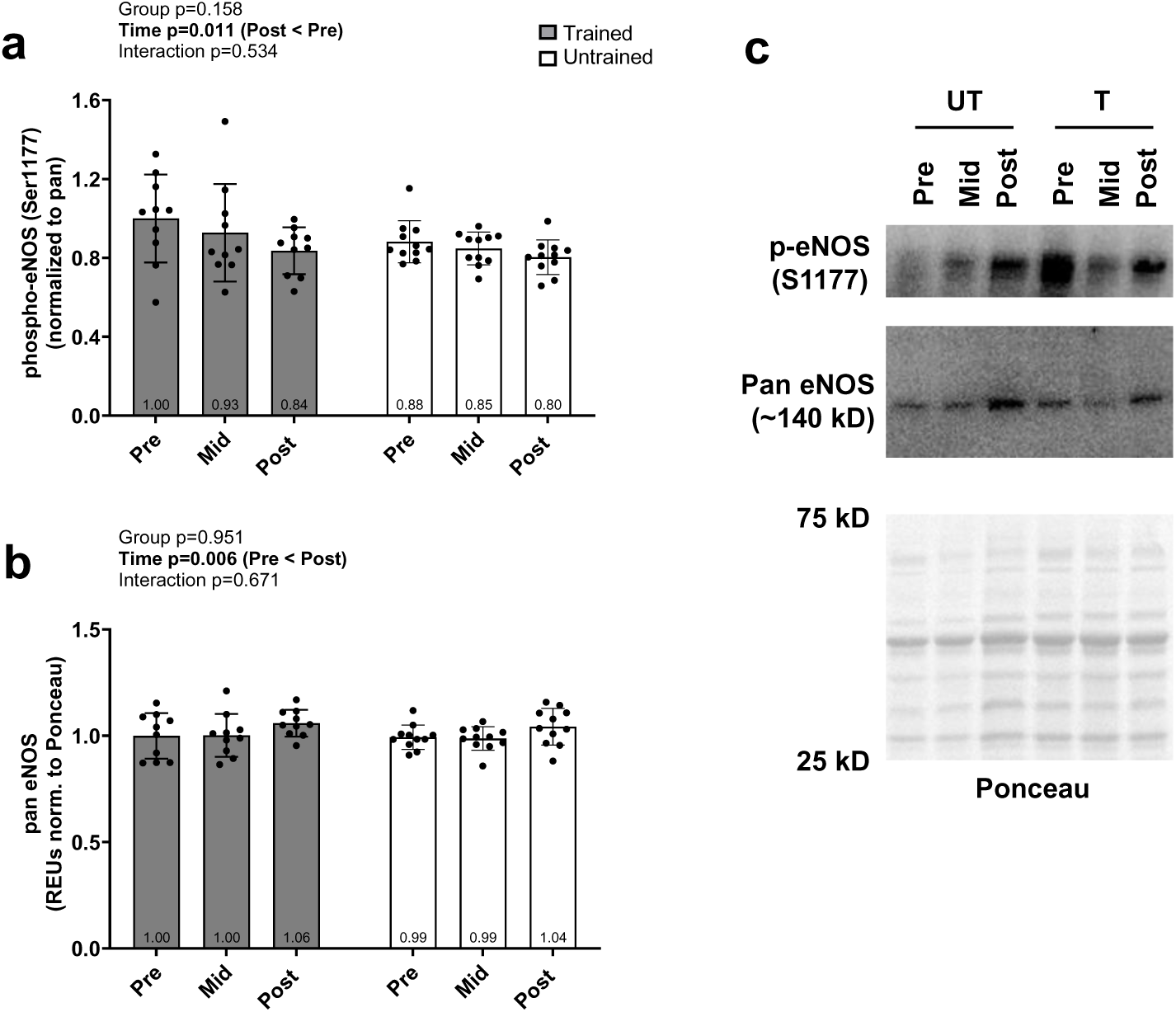
Phosphorylated and pan eNOS responses. Data presented as mean ± standard deviation for phosphorylated-eNOS (a) and pan protein levels (b) prior to the intervention (Pre), following two weeks of leg disuse (Mid), and following 8 weeks of recovery resistance training (Post). Panel c shows representative western blots. Abbreviations: T, trained participant; UT, untrained participant. Other note: each participant’s data point is overlain on bar graphs and mean values are depicted at bottom of bar.

### Changes in capillaries per fiber

Figure 4 contains skeletal muscle type I and II myofiber capillary count data. Type I capillary number did not exhibit significant main or interaction effects (Fig. 4a). However, Type II capillary number did exhibit a significant main effect of time whereby POST was greater than PRE in all participants (p=0.014, Fig. 4b).

**Figure 4.**
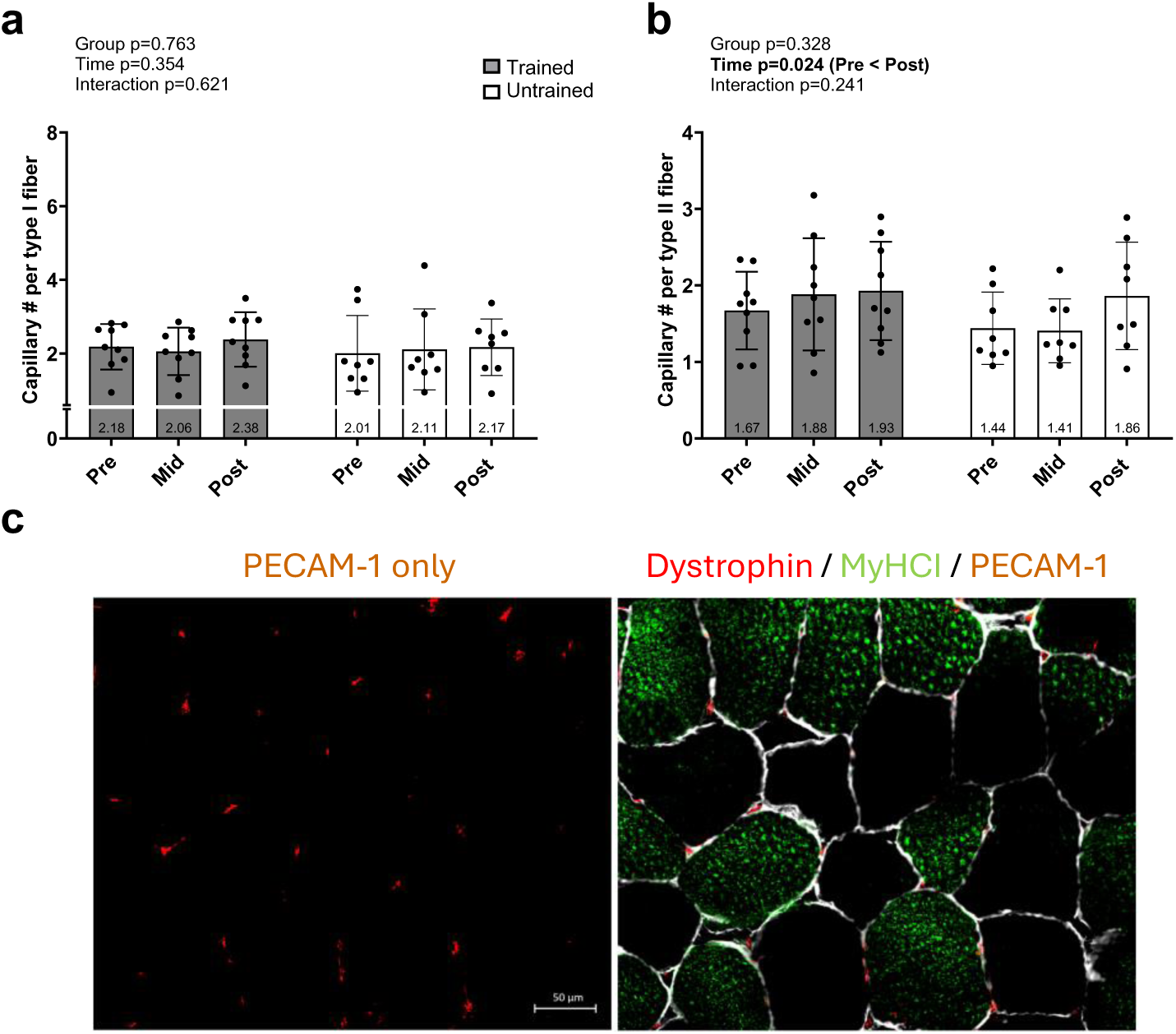
Type I and II myofiber capillary responses. Data presented as mean ± standard deviation for type I (a) and type II (b) myofiber capillary number prior to the intervention (Pre), following two weeks of leg disuse (Mid), and following 8 weeks of recovery resistance training (Post). Panel c shows a representative 20x fluorescent image of PECAM-1 only and a merged image; note, dystrophin (Cy5) is pseudocolored white. Abbreviations: T, trained participant; UT, untrained participant. Other note: each participant’s data point is overlain on bar graphs and mean values are depicted at bottom of bar.

### Associations between changes in type II fiber capillary number and myofiber size

Given that type II fiber capillary number increased with RT regardless of training group, we were interested in determining if this outcome was associated with percentage changes in type II fiber cross-sectional area (fCSA) values. Michel et al. reported type II fCSA values in 21 participants, and the 16 participants assayed herein demonstrated numerical decreases from PRE to MID (6225±1593 to 5943±1566 µm^2^, p=0.137) as well as values at POST (7165±1418 µm^2^) that were greater than PRE (p=0.019) and MID (p<0.001). However, no significant correlations existed for percentage changes in type II fiber capillary number and type II fCSA from PRE-to-MID (r= 0.020, p=0.941), MID-to-POST (r= 0.392, p=0.133), or PRE-to-POST (r= −0.120, p=0.657).

## DISCUSSION

Reduced skeletal muscle angiogenesis and losses in capillary content (i.e., capillary rarefaction) have been posited to play a role in skeletal muscle disuse-induced atrophy (31, 32). Moreover, various skeletal muscle atrophy models in rodents suggest capillary rarefaction and/or suppressed angiogenesis coincide with skeletal muscle atrophy (25, 33–38). Various rodent and human studies have sought to assess how reloading following disuse atrophy effects various markers related to angiogenesis (39–42). This is highlighted in that evidence in human participants suggests that 4-6 weeks of endurance training following two weeks of leg immobilization alters markers suggestive of increased angiogenesis (27, 43). However, no human study to date has examined how recovery RT following disuse atrophy affects these outcomes. Therefore, this study aimed to assess alterations in markers associated with skeletal muscle angiogenesis following two weeks of leg immobilization immediately followed by 8 weeks of subsequent recovery RT. In agreement with our hypotheses, 2 weeks of leg immobilization is seemingly anti-angiogenic as evidenced through a decrease in VEGF and increase in TSP-1 protein levels. Also, in agreement with our hypotheses, 8 weeks of recovery RT enhances angiogenesis as evidenced through MID-to-POST increases in VEGF and VEGFR2 protein levels as well as a PRE-to-POST increase in type II fiber capillary number. Prior to a more expanded discussion of the implications of our findings, it is worth noting that training history differences in most outcomes were not evident. This finding also agrees with the parent publication by Michel et al. (28) who reported that atrophic and hypertrophic responses, along with corresponding molecular targets, were similar regardless of training status/history in the given model.

Extracellular matrix remodeling markers associated with angiogenesis remained relatively unchanged with disuse and subsequent recovery RT, albeit TIMP1 protein levels were responsive across time points. Several factors directly related to angiogenesis (VEGF, VEGFR2, and TSP-1) were dynamically altered following 2 weeks of disuse. TSP-1 is a potent inhibitor of angiogenesis by antagonizing VEGF and reducing endothelial cell proliferation and migration (19, 44). In rodent models, skeletal muscle capillary number is reduced with the administration of a TSP-1 mimetic (45), and conversely, capillary number is elevated in a global TSP-1 knockout model (46). Interestingly, our findings that two weeks of leg immobilization reduce VEGF and increase TSP-1 protein levels in skeletal muscle agree with another human study demonstrating similar effects following two weeks of leg bracing (43). However, in the current model of disuse, we did not observe capillary rarefaction despite decreases in skeletal muscle VEGF and VEGFR2 as well as an increase in TSP-1. Our VEGF, VEGFR2 and TSP-1 data suggest that observed protein expression patterns precede marked changes in (19) and that skeletal muscle capillary rarefaction may be observed with a longer duration or with a more severe disuse model/stimulus.

Resistance training consistently leads to skeletal muscle hypertrophy, and this has been shown to coincide with increased skeletal muscle capillarization (5, 7, 47–52) along with acute upregulations of angiogenic markers after a single bout or short-term RT (5, 13–15, 53), Furthermore, eNOS (an enzyme known to promote vasodilation and angiogenesis) is also altered with exercise (16, 19). In the current study, we report an increase in VEGF, VEGFR2, and eNOS protein levels from MID to POST time points. This suggests dynamic interplay between angiogenic markers and following disuse atrophy and subsequent recovery RT. The observed increases in type II myofiber capillary number agree with other human studies reporting that RT increases skeletal muscle capillary number (5, 48, 49, 54). Despite increases in capillary number, no associations were shown between percentage changes in fCSA and capillary number in the current study. In contrast with our data, skeletal muscle capillarization has been shown to be predictive of the skeletal muscle hypertrophic response to RT among older adults (8, 9). While speculative, our lack of associations may be due to younger/healthy adults being investigated compared to the studies previously mentioned. In the current model, several markers known to regulate angiogenesis are dynamically altered following both disuse and recovery RT, while protein levels of these markers are affected, capillary number is only significantly altered in type II myofibers following RT and is not associated with myofiber hypertrophy. Taken together, the data suggest that angiogenic-related markers are altered in both an atrophic and hypertrophy stimulus but do not seem to drive changes in skeletal muscle mass in a young healthy population. Further interrogations comparing different aging populations following both disuse and RT stimulus are needed to fully unveil any further associations between skeletal muscle capillarization and skeletal muscle hypertrophic adaptations.

### Limitations

Similar to many human participant RT studies, this study is limited by a small sample size. A larger sample size with a more diverse participant pool would likely increase the generalizability of our findings. Moreover, the participants herein were predominantly males who underwent non-complicated disuse whereby the braced limb was not injured, and the immobilization period was among a less severe model of immobilization (e.g., bedrest, limb casting, microgravity). Finally, biopsies were only performed in the basal state, so we did not interrogate markers of gene regulation (e.g., mRNA expression) related to angiogenesis that might be acutely responsive to RT and/or disuse atrophy. Indeed, these markers would add mechanistic insight to how the model may affect phenotypic outcomes.

### Conclusions

This study furthers findings from our original study by Michel et al. (28) by providing the first human evidence demonstrating that leg immobilization and subsequent recovery RT dynamically alter skeletal muscle markers associated with angiogenesis in younger adults. However, further investigation is needed to determine if our findings are consistent across other disuse models (e.g., complicated disuse with casting or bedrest) and different participant cohorts (e.g., older individuals).

## ADDITIONAL INFORMATION

## ACKNOWLEDGEMENTS

We would like to acknowledge the participants who completed this study.

## DATA AVAILABILITY STATEMENT

Data that support the findings of this study are available from the corresponding author upon reasonable request.

## CLINICAL TRIAL REGISTRATION

This study was registered as a clinical trial at clinicaltrials.gov (NCT05760066).

## ETHICS APPROVAL STATEMENT

Study protocols were carried out in accordance with the most recent version of the declaration of Helsinki. All study procedures were approved by Auburn University’s Institutional Review Board (approval number: 23-220 MR 2305).

## PATIENT CONSENT STATEMENT

All participants in this study provided verbal and written consent in accordance with the above IRB approval.

## AUTHOR CONTRIBUTIONS

MCM and MDR primarily drafted the manuscript and prepared figures. HSB is a licensed physical therapist who assisted with oversight throughout the intervention. MSS provided critical equipment and expertise for the carrying out of the leg immobilization protocol. MCM primarily carried out laboratory-based assays. JMM, JSG, DLP, DAA, MLM, AAB, NJK, and CBM provided crucial assistance with original data collection procedures and/or consistent input for the duration of this follow-up investigation. All co-authors assisted with revising and editing the manuscript, and all co-authors approved the final version.

